## Supplemental Figures and Tables for "Deciphering DEL Pocket Patterns through Contrastive Learning"

### **Supporting Information for**
### Deciphering DEL Pocket Patterns Through Contrastive Learning


Wenyi Zhang[1,2], Yuxing Wang[1,2], Rui Zhan[1,2], Runtong Qian[1,2], Qi Hu[1,2], Jing Huang[1,2,*].

1. State Key Laboratory of Gene Expression, School of Life Sciences, Westlake University, 600 Dunyu Road, Hangzhou, Zhejiang 310030, China.

2. Westlake AI Therapeutics Lab, Westlake Laboratory of Life Sciences and Biomedicine, 18 Shilongshan Road, Hangzhou, Zhejiang 310024, China.

*Corresponding author here


**This PDF file includes:**

Figures S1 to S19
Tables S1 to S2

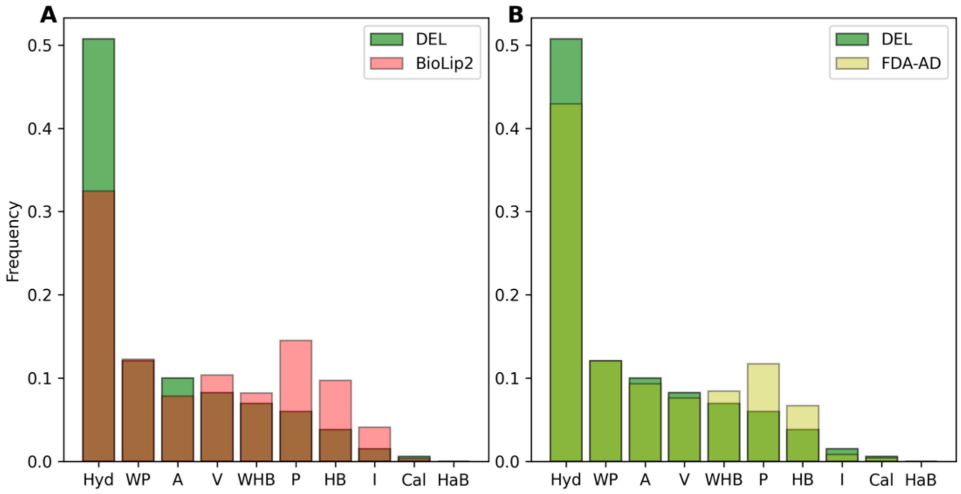


**Figure S1. Comparison of 10 different types of interactions between pockets and ligands in the DEL, BioLiP2, and FDA-AD datasets.** The interactions are represented by the following types: 'V' (Van der Waals), 'HB' (Hydrogen Bond), 'WHB' (Weak Hydrogen Bond), 'HaB' (Halogen Bond), 'I' (**I**onic), 'A' (Aromatic), 'Hyd' (Hydrophobic), 'Cal' (Carbonyl), 'P' (Polar), and 'WP' (Weak Polar).

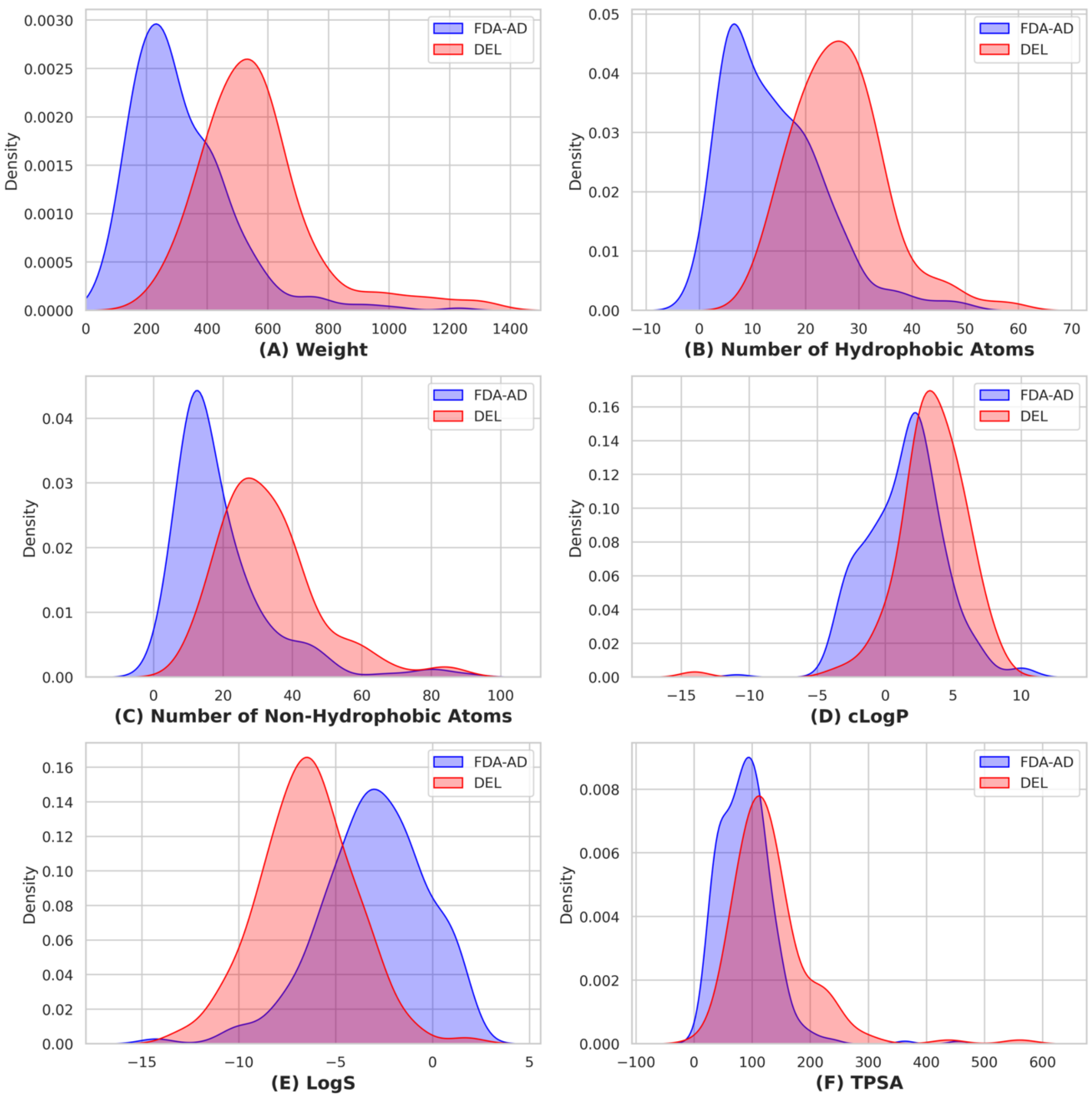


**Figure S2. Comparison of physicochemical properties between DEL molecules and FDA approved drugs.** The properties analyzed include molecular weight, the number of hydrophobic atoms, the number of hydrogen atoms, the log octanol/water partition coefficient (cLogP), the log solubility in water (logS), and the topological polar surface area (TPSA).

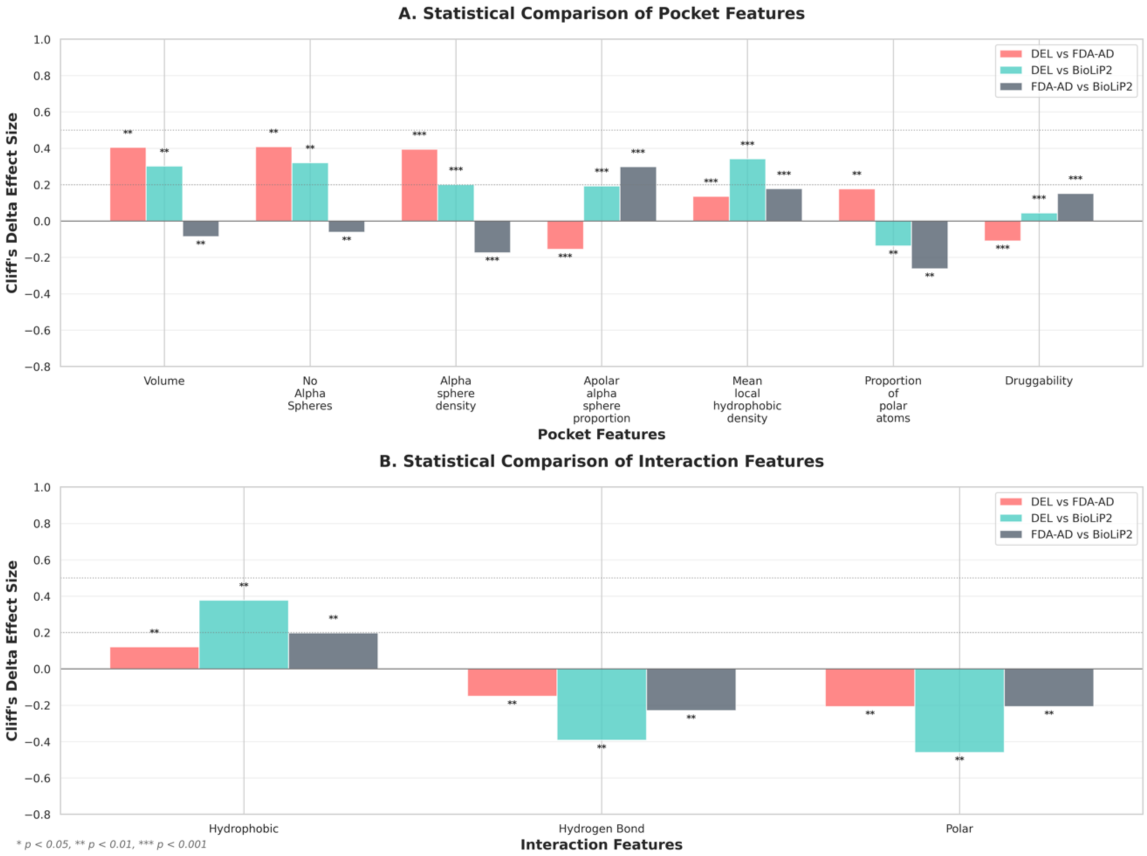


**Figure S3. Statistical comparison of pocket and interaction features. A.** Pocket features (volume, alpha spheres, density, polar/apolar composition, hydrophobicity, druggability) compared between DEL, FDA-AD, and BioLiP2 targets using Cliff's delta; significance: * $p < 0.05$, ** $p < 0.01$, *** $p < 0.001$; **B.** Interaction features (hydrophobic, hydrogen-bond, and polar contacts) compared across the three datasets using Cliff's delta with the same significance notation.

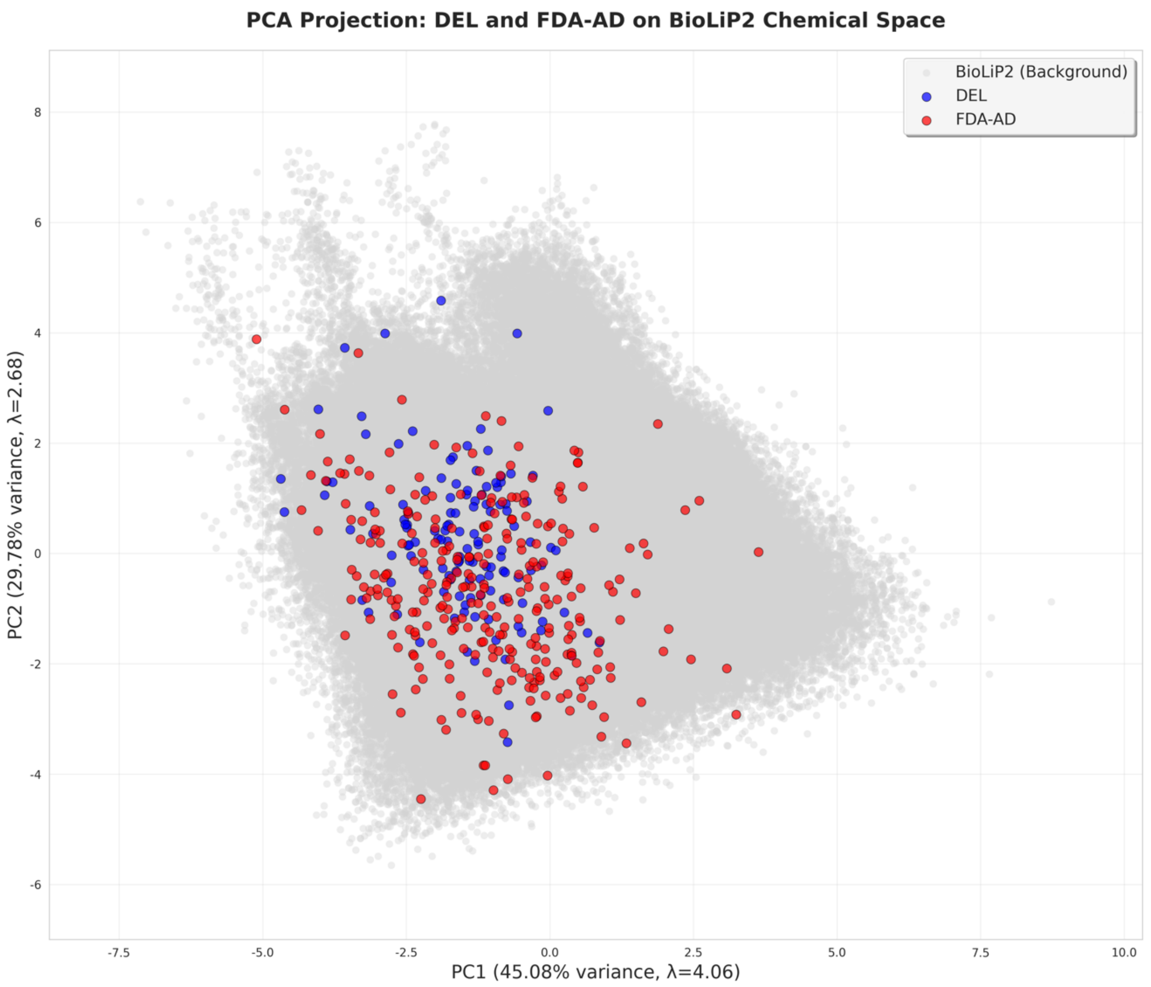


**Figure S4. Principal component analysis (PCA) of pocket and interaction features.** DEL (blue) and FDA-AD (red) targets were projected onto the PCA space trained using BioLiP2 pockets (gray background). Kernel density contours highlight the distinct yet partially overlapping distributions of DEL and FDA-AD targets. PC1 is primarily influenced by chemical composition, including apolar–polar balance and interaction types (e.g., apolar α-sphere proportion 4.1%, proportion of polar atoms 4.0%, hydrophobic 3.8%, hydrogen bonds 3.9%, polar interactions 4.0%), whereas PC2 is dominated by structural size descriptors (pocket volume 11.7% and number of alpha spheres 11.2%). Together, these components capture the separation of DEL targets from reference datasets.

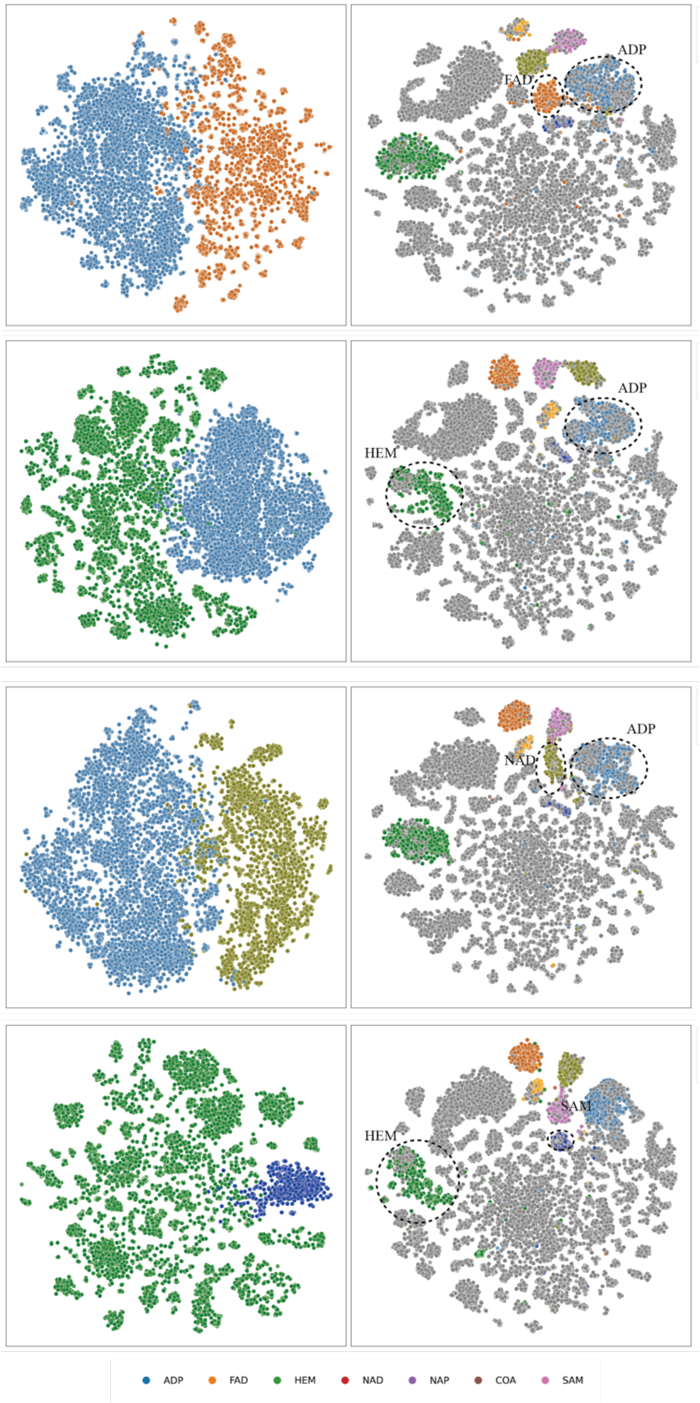


**Figure S5**. **t-SNE visualization of the model's performance in classifying excluded binding pocket types during the ablation study.** The study involved excluding different combinations of binding pockets from the BioLiP2 training dataset: ADP and NAD pockets (A and B), HEM and ADP pockets (C and D), ADP and NAD pockets (E and F), and HEM and SAM pockets (G and H).

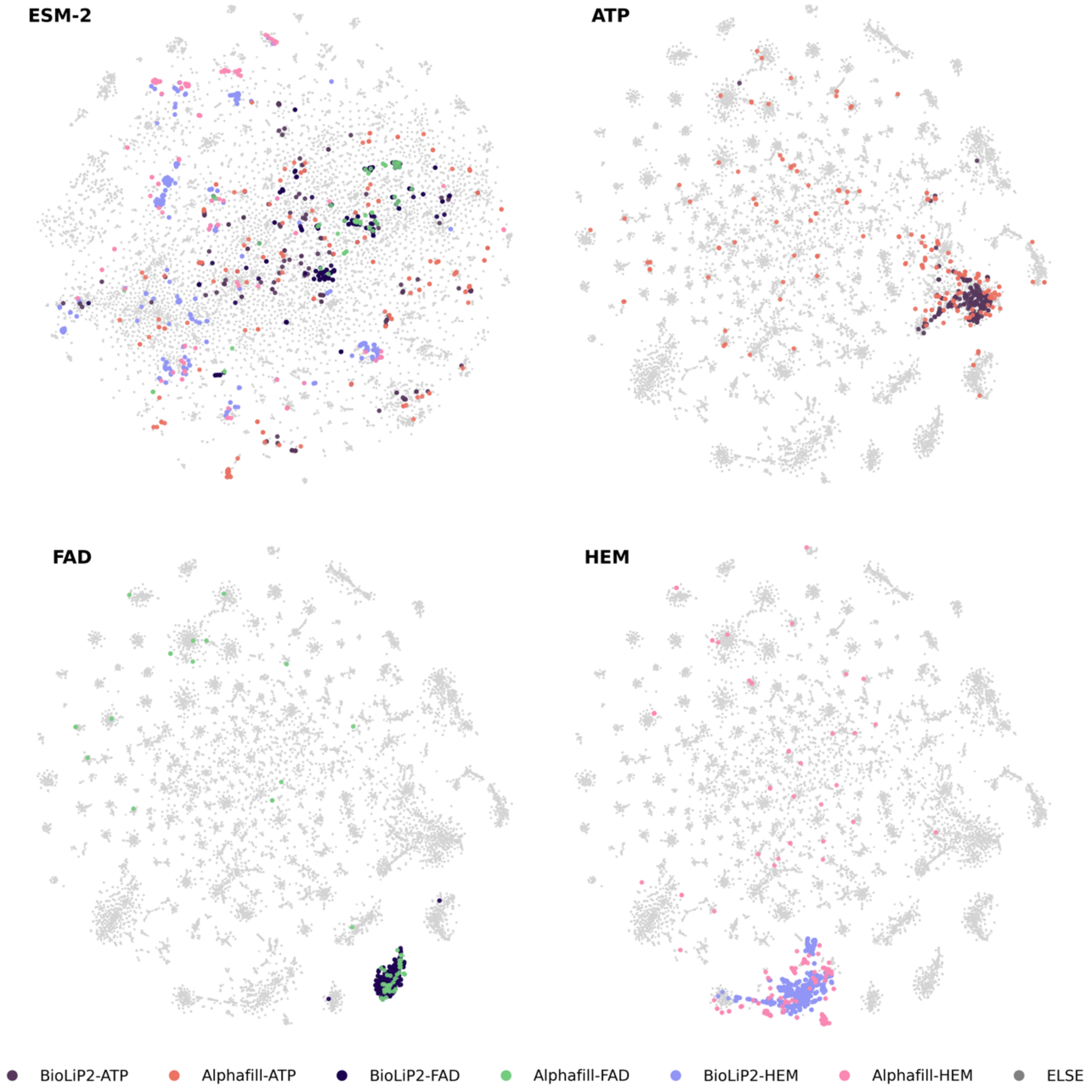


**Figure S6. t-SNE clustering of ATP, FAD, and HEM binding pockets from BioLiP2 and Alphafill datasets.** (**A**) Clustering with ESM2 features. (**B**) ErePOC clustering for ATP pockets. (**C**) ErePOC clustering for FAD pockets. (**D**) ErePOC clustering for HEM pockets.

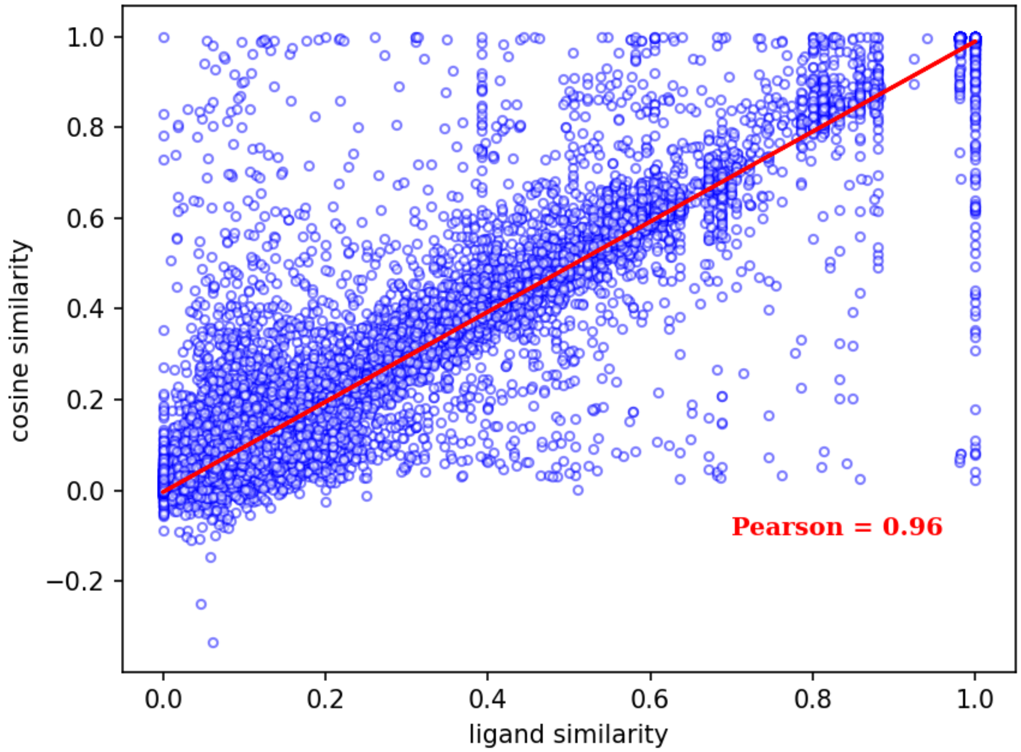


**Figure S7. Pearson Correlation Analysis between ligand Tanimoto similarity and pocket cosine similarity derived from the ErePOC vector**.

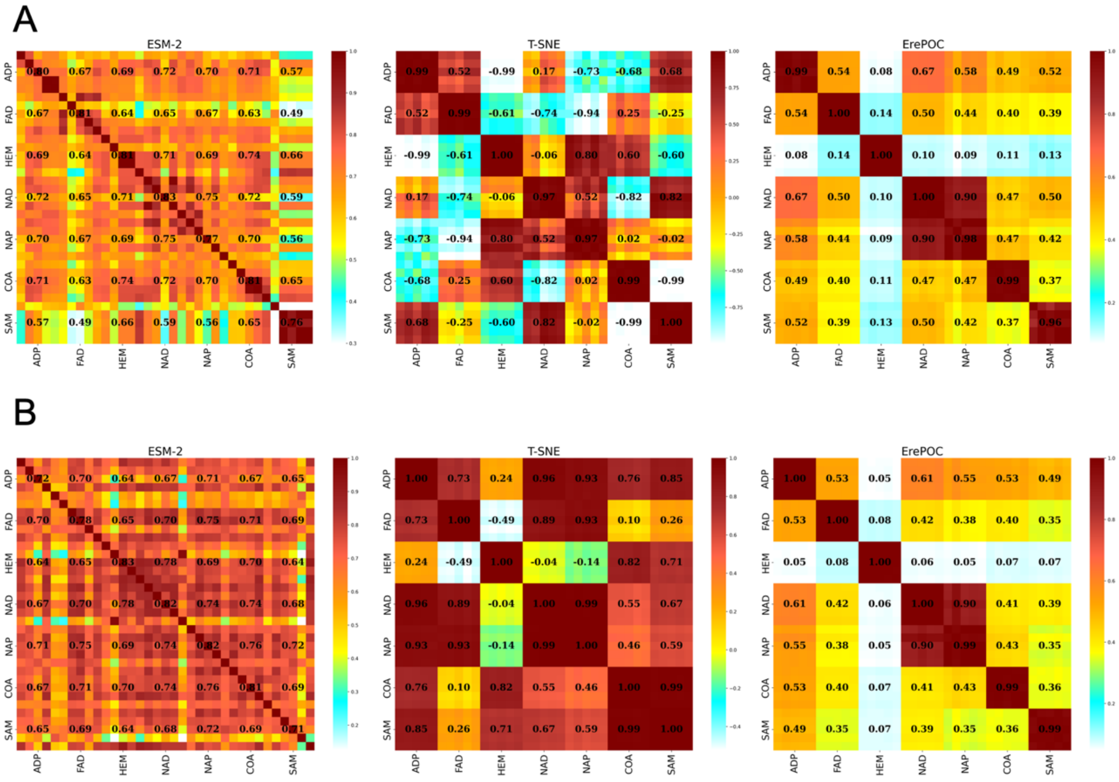


**Figure S8. Cosine similarity comparison across different binding pockets under two BioLiP2 pocket definitions. A.** Cosine similarity results based on the original BioLiP2 annotation–defined pockets, including ESM-2 vectors, t-SNE 2D representations after ErePOC transformation, and ErePOC embeddings; **B.** Corresponding results obtained using the unified 5 Å pocket definition. The overall similarity distributions remain consistent across the two definitions, confirming the robustness of ErePOC embeddings with respect to pocket boundary choice.

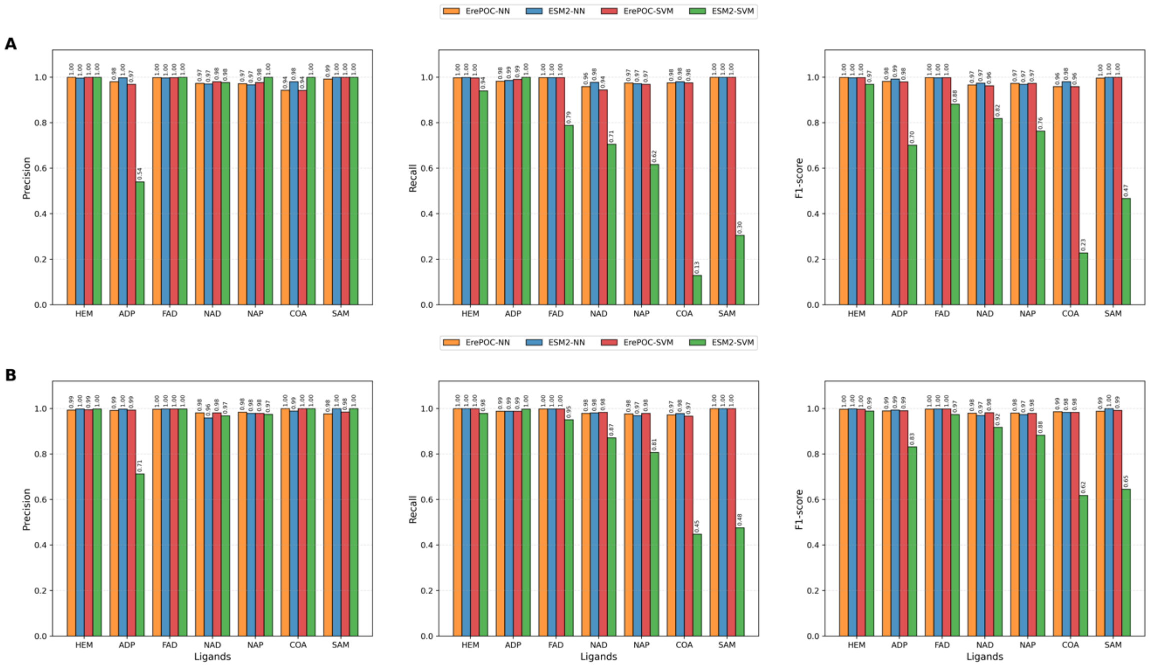


**Figure S9. Comparative performance of four models under two BioLiP2 pocket definitions. A.** Predictive performance of ErePOC-NN, ErePOC-SVM, ESM2-NN, and ESM2-SVM using BioLiP2 annotation–defined pockets; **B.** Corresponding results using the unified 5 Å pocket definition.

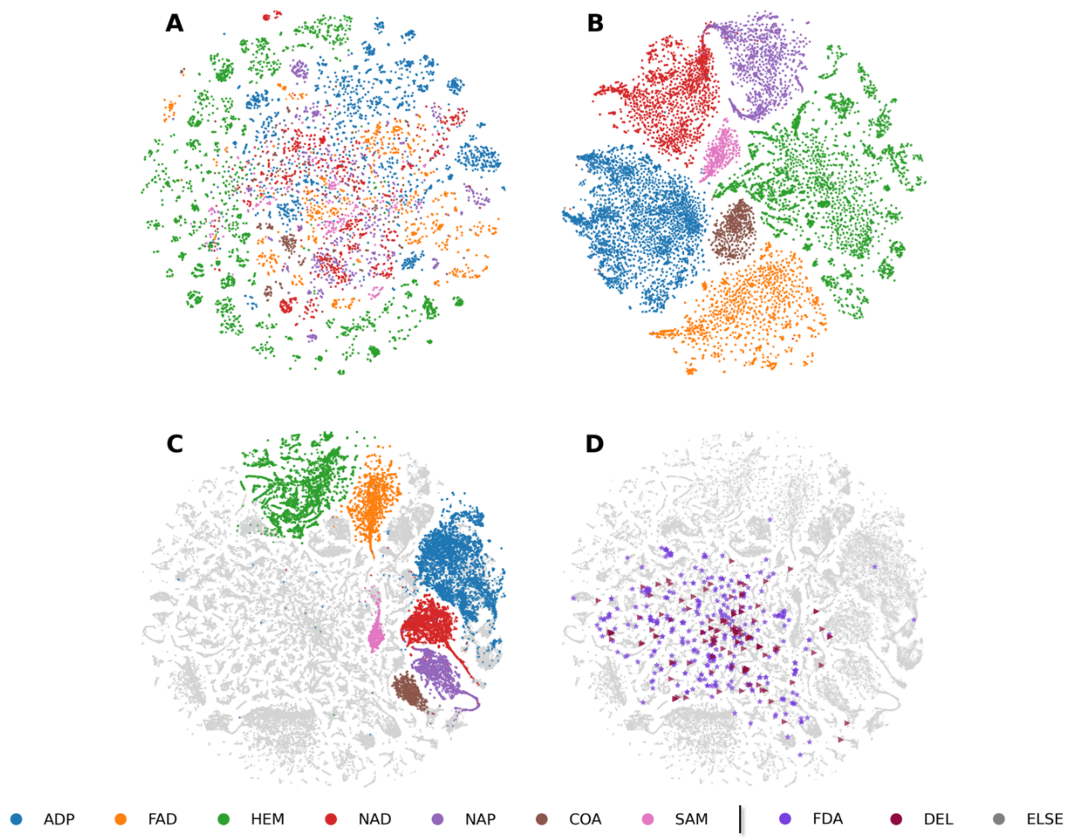


**Figure S10. t-SNE visualization of ErePOC and ESM-2 representations for BioLiP2 dataset using the 5 Å pocket definition. A–B.** Visualization of the seven types of ligand-binding pocket landscapes using ESM-2 (A) and ErePOC (B) representations, respectively. **C.** Pocket landscape using ErePOC representations. **D.** Comparison of pocket landscapes for the FDA-AD and DEL datasets against BioLiP2 using ErePOC representations. Results are consistent with those obtained using curated BioLiP2 binding annotations (Figure 4), confirming the robustness of the ErePOC embedding with respect to pocket definition.

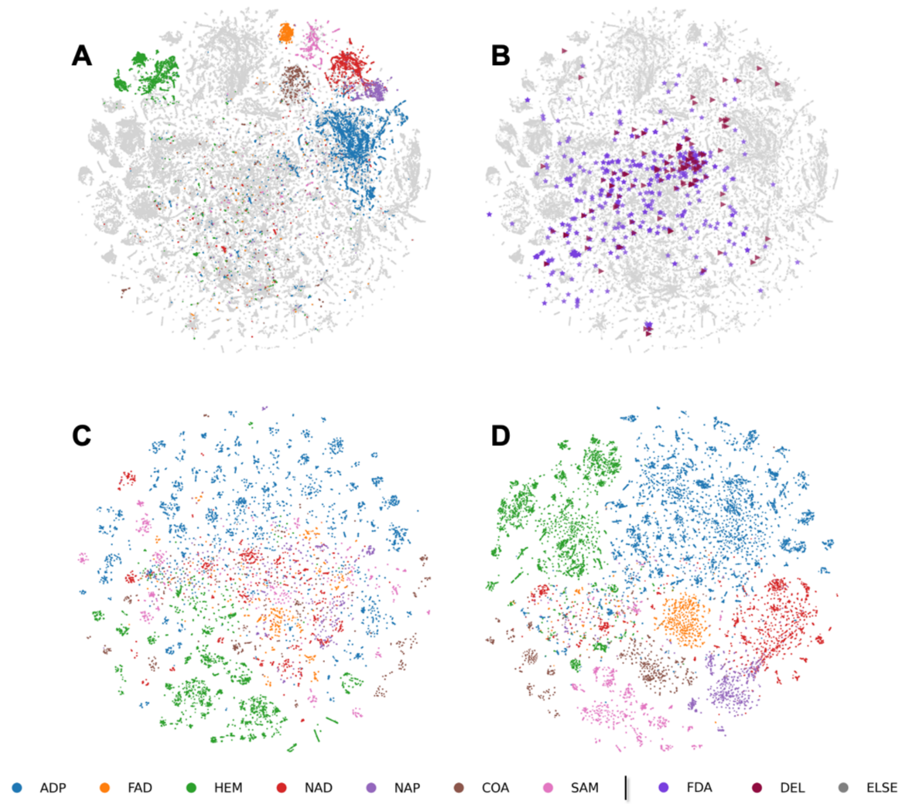


**Figure S11. t-SNE visualization of ErePOC and ESM2 representations for AlphaFill dataset**, where ErePOC was trained on BioLiP2 annotation–defined pockets. **A.** Pocket landscape using ErePOC representations. **B.** Comparison of pocket landscapes for the FDA-AD and DEL datasets against AlphaFill using ErePOC representations. **C-D.** Visualization of the 7 types of ligand-binding pocket landscapes using ESM2 (C) and ErePOC (D) representations, respectively.

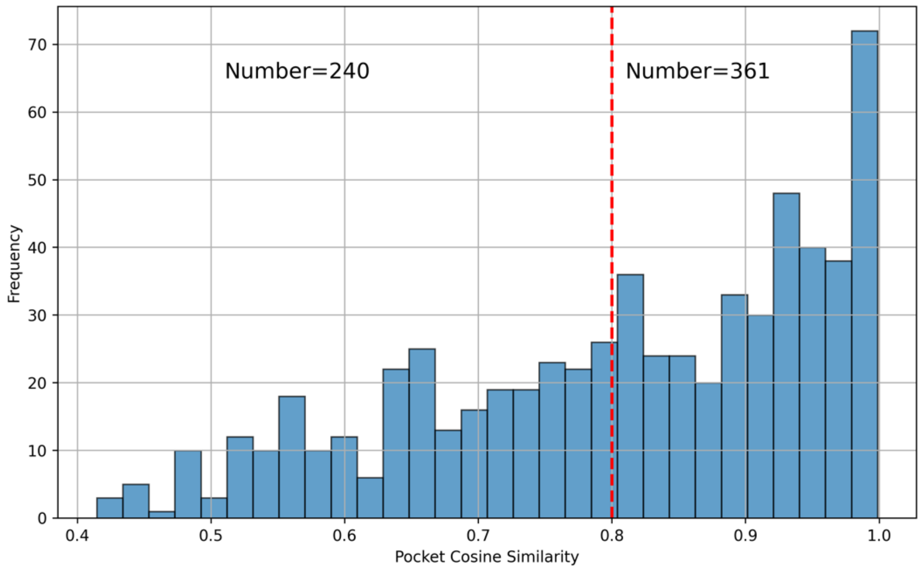


**Figure S12**. Distribution of DEL neighbor cosine similarity scores in BioLiP2.

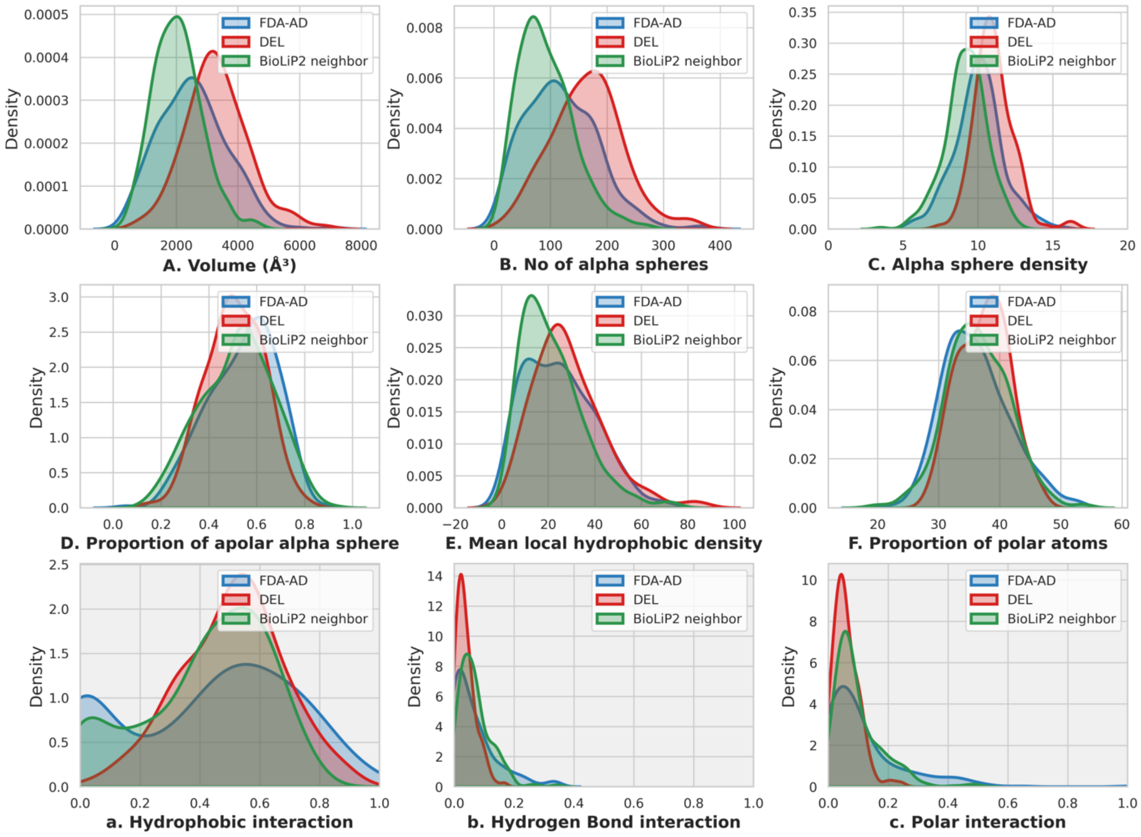


**Figure S13**. Physicochemical properties of pockets and ligand-pocket interaction analysis for DEL neighbors.

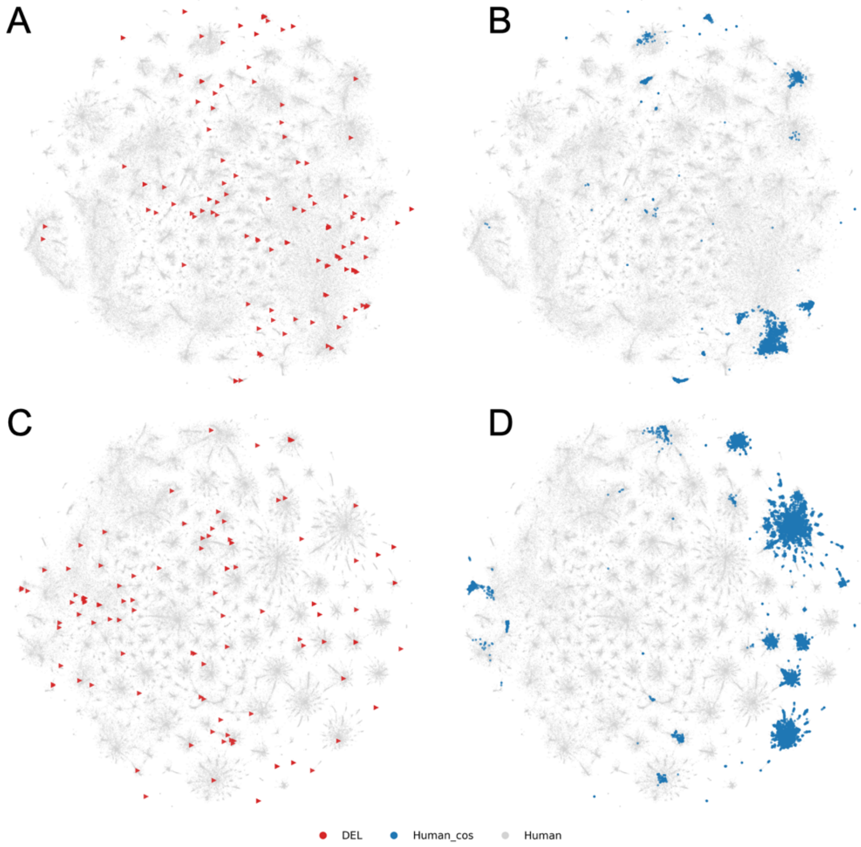


**Figure S14. t-SNE visualization of DEL pockets and human pockets with cosine similarity > 0.8. A-B.** Results obtained using the model trained on BioLiP2 annotation–defined pockets. **C-D.** Results obtained using the model trained with pockets defined as all residues within 5 Å of the bound ligand.

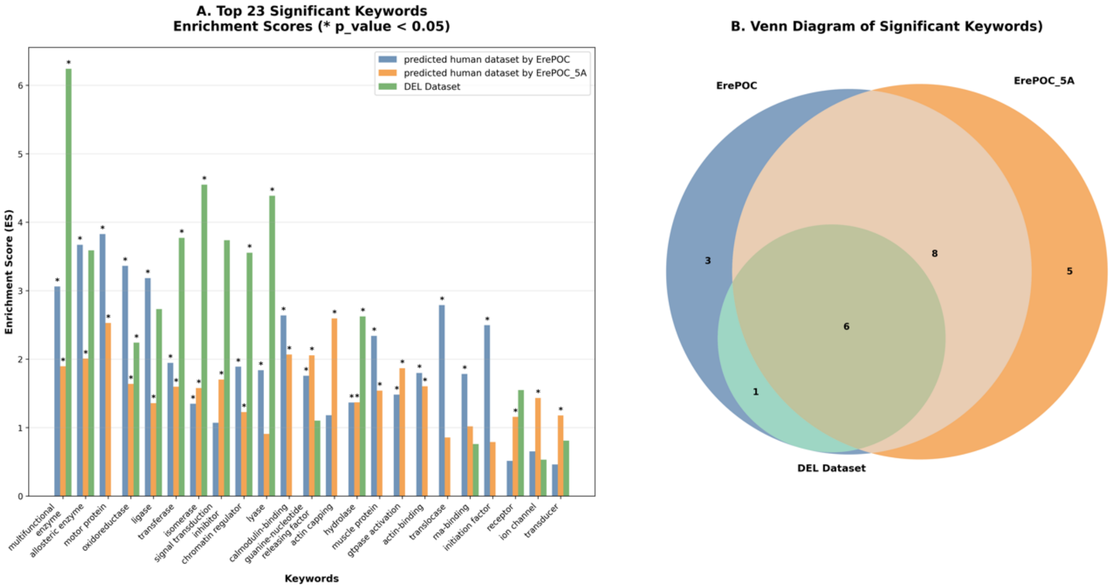


**Figure S15. Comparative analysis of significantly enriched keywords across predicted human proteome datasets and DEL dataset. A.** Top 23 significantly enriched keywords showing enrichment scores (ES) for predicted human dataset by ErePOC (blue), predicted human dataset by ErePOC_5A (orange), and DEL dataset (green). Asterisks (*) indicate statistical significance (p-value < 0.05); **B.** Venn diagram illustrating the overlap of significantly enriched keywords among the three datasets, demonstrating the shared and unique functional features identified by each prediction method. ErePOC_5A means the model was trained using the 5 Å pocket definition.

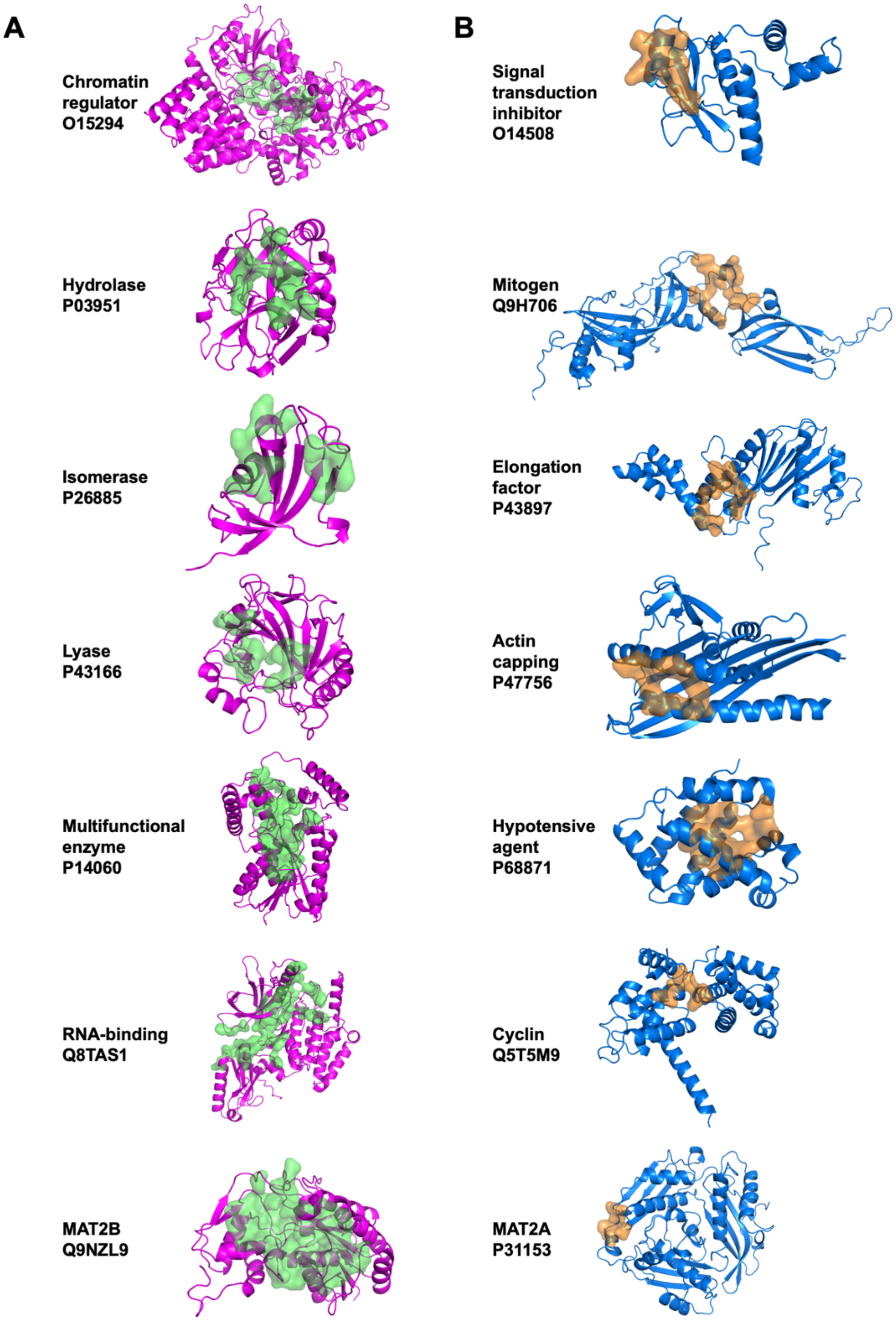


**Figure S16. Selected human proteins and their pockets for in silico DEL screening.** This figure shows the 14 human protein targets selected for large-scale virtual screening on a DEL compound library, grouped into DEL-enriched (left panels) and DEL-neutral (right panels) categories based on computational predictions. The MAT2B and MAT2A (bottom panels) are included as an example for different predicted DEL compatibility within a single protein family.

These classifications support the experimental design used to evaluate ErePOC-predicted DEL suitability.

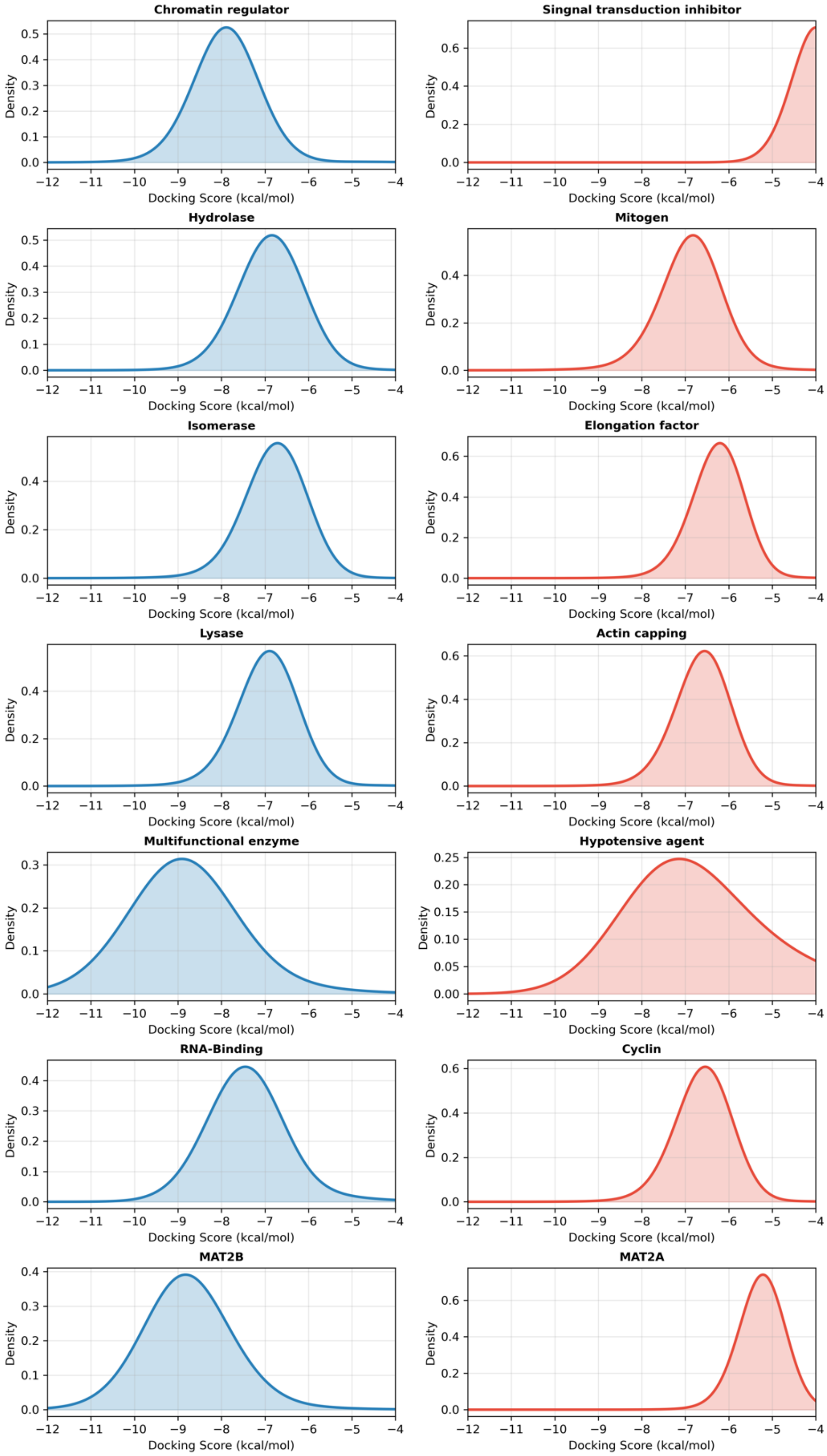


**Figure S17. Docking score distributions for the 14 human protein targets.**

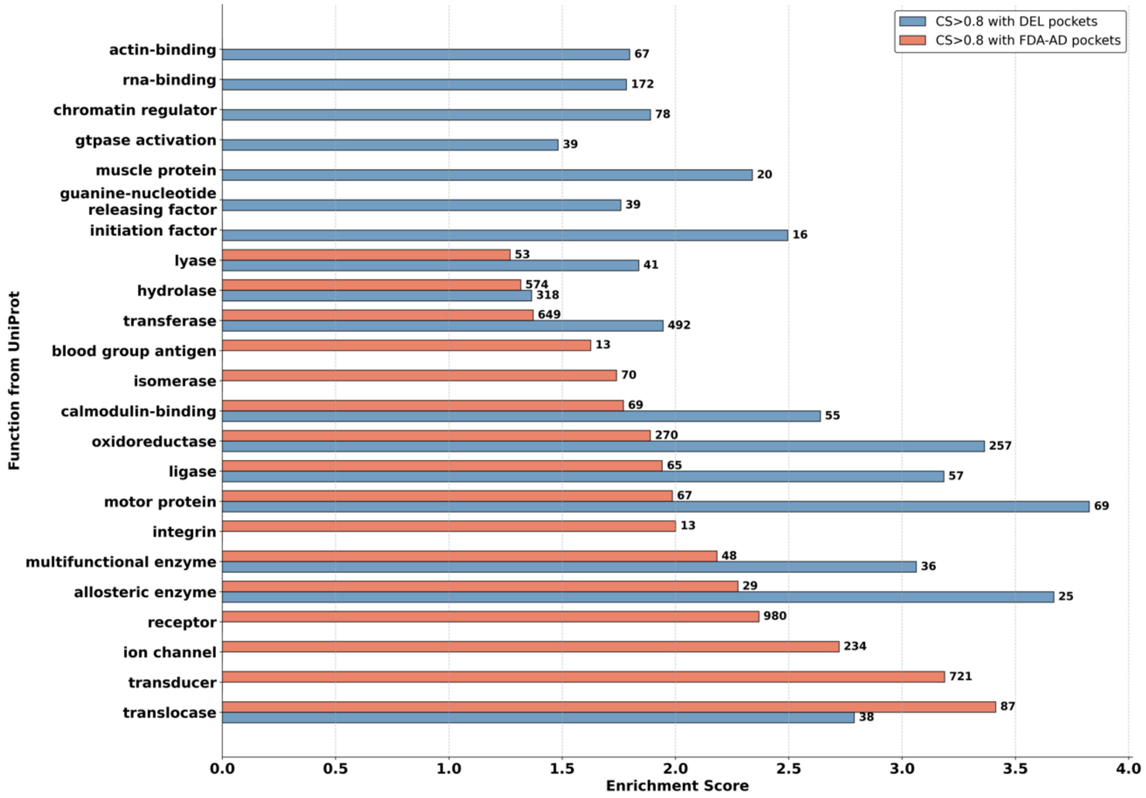


**Figure S18. Comparative analysis of human protein enrichment using pocket similarity with either DEL or FDA-AD pockets using ErePOC representation (cosine similarity > 0.8)**

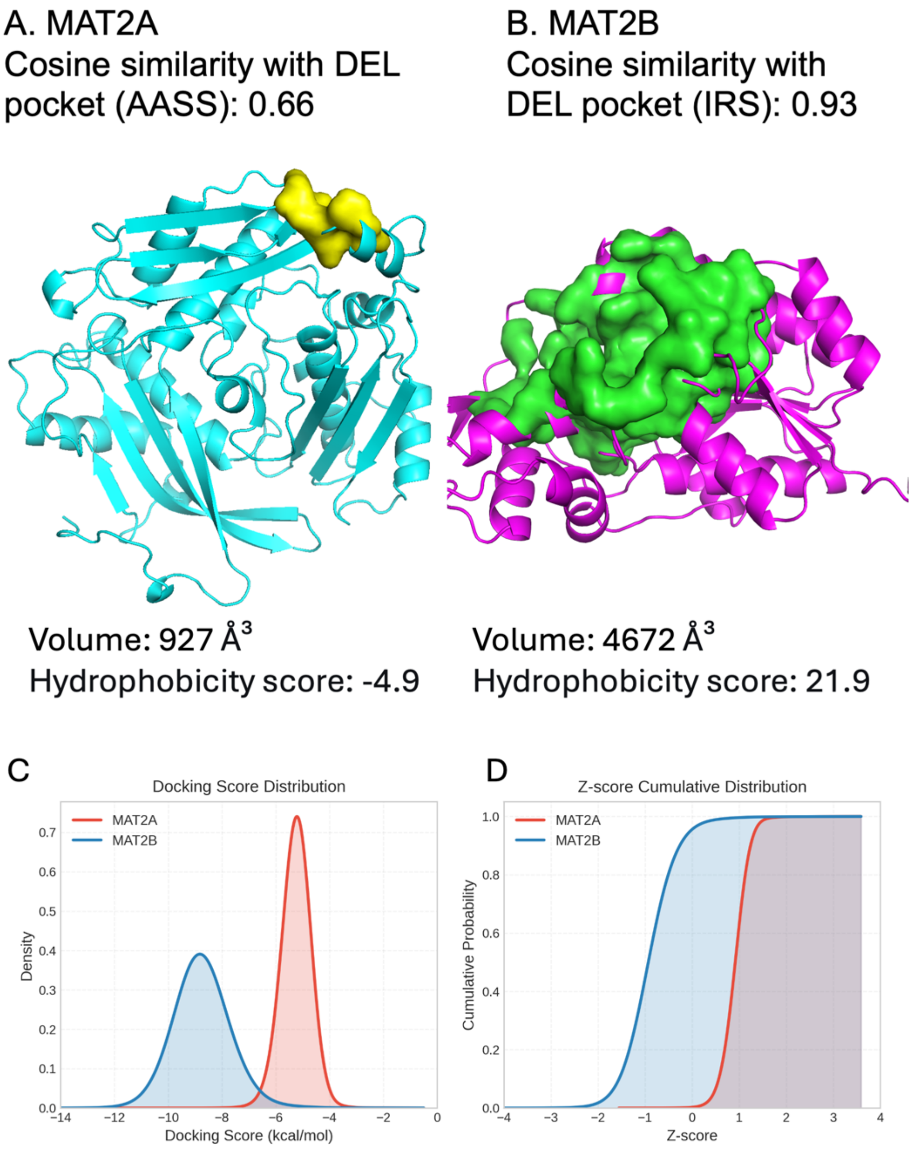


**Figure S19. Comparative Analysis of Pocket Properties between MAT2A and MAT2B. A.** The structure and pocket of MAT2A; **B.** The structure and pocket of MAT2B; **C.** Distribution of docking scores; **D.** Distribution of docking Z-scores for MAT2A (blue) and MAT2B (red).

**Table S1. Summary of datasets included in the study**

| Dataset | Size |
|---|---|
| DEL | 128 |
| FDA-AD | 340 |
| BioLiP2 | 326,416 |
| Alphafill | 293,019 |
| Homo sapiens species | 23,391 |

BioLiP2 dataset: Multiple structures bound to the same ligand are retained intentionally, allowing ErePOC to learn from diverse conformational states of the same ligand across different proteins and crystallization conditions. This enhances functional relevance and improves generalizability.

AlphaFill dataset: Similarly, AlphaFill contains redundant ligand–protein complexes. Redundancy was maintained to expose the model to a broad spectrum of protein contexts and ligand environments. After filtering for steric clashes, 293,019 high-confidence complexes were retained.

FDA-AD dataset: Unique crystal structures were defined as unique FDA drug–protein pairs. Redundant structures for the same drug–target pair were removed, but multiple targets of the same drug are retained.

DEL dataset: A one-to-one mapping was applied by retaining only the highest-affinity DEL hit per target, resulting in 128 unique DEL–target structures.

**Table S2. Detailed information of the DEL target protein structures dataset**

| Index | Target | PDB ID | SMILE for DEL compound |
|---|---|---|---|
| 1 | 3CLpro | 7vu6A | Brc1cc([N+](=O)[O-])c(NC2C(NC(=O)c3c4c(cnc3)cccc4)CCCC2)c(C(=O)NC)c1 |
| 2 | 53BP1 TTD | 4x34A | O=C(N)C(NC(=O)C=1C(c2c(CNC(=O)c3nccc(N4CC[NH+](C)CC4)c3)cccc2)CC=CC=1)CCCC[NH+]1CCCC1 |
| 3 | AASS | 5l78A | Clc1cnc(OC2CC(NC(=O)C(NC(=O)c3c(SCCS(=O)(=O)c4ccccc4)nccc3)CCC(=O)NC(C(=O)O)CC(=O)N)CCC2)cc1 |
| 4 | ADAMTS-4 | 4wk7A | O=C(N(CC)c1cc(C)ccc1)c1ccc(CNc2nc(NC[NH+]3CC[NH+](C)CC3)nc(N3Cc4c(cc(OC)c(OC)c4)CC3)n2)cc1 |
| 5 | AGP | 3kq0A | Clc1c(C2NC(=O)C(CCN=[N+]=N)NC(=O)C(C)Cc3c(C(=O)NCCNC(=O)C2)cccc3)cccc1 |
| 6 | AGP; CA IX | 6rzxA | S(=O)(=O)(N(CC(=O)NCCOCCC(=O)NCCOCCC(=O)NC(C(=O)[O-])CCCCNC(=O)C=Cc1oc(-c2c(C(F)(F)F)cccc2)cc1)c1c(-c2ccccc2)cccc1)c1ccc(C)cc1 |
| 7 | AT1R | 4zudA | Fc1ccc(CC(NC(=O)C(NC(=O)C(NC(=O)C=Cc2cc(O)c(O)cc2)C)CC2CCCCC2)C(=O)OCC)cc1 |

| | | | |
|---|---|---|---|
| 8 | ATAD2 | 6hi4A | Clc1c(C(=O)NC(C[NH2+]C2CCC([NH3+])CC2)Cc2ccc(C#N)cc2)cc(c(OC)c1)-c1oc(C[NH2+]C(C)c2ccc(C)cc2)cc1 |
| 9 | ATX | 4zgaA | S(=O)(=O)(C)c1ccc(NC2(C)CCN(C(=O)C(NC(=O)c3c(F)ccc(CC)c3)C(C)C)CC2)cc1 |
| 10 | Aurora A | 2xngA | O=C(O)Cn1nnc(C[NH+]2Cc3c(ccc(-c4ccc(C=O)cc4)c3)C2)c1 |
| 11 | autotaxin | 6w35A | FC(F)(F)c1cc(C(=O)NC(C(=O)N2CCC3(N(c4cc5c([nH]nc5)cc4)C(=O)N(C)C3=O)CC2)C(C)C)c(F)cc1 |
| 12 | β2AR | 5x7dA | Fc1ccc(CC(NC(=O)C(NC(=O)C2(NC(=O)C=Cc3cc(O)c(O)cc3)CC2)CC2CCCCC2)C(=O)OCC)cc1 |
| 13 | β2AR; CysLT | 7dhrR | O=C(NCCc1ccc(O)cc1)C1(C)CC2C(C)(CCC3(CC)C4(C)C(C(=O)C=C23)C2(C)C(C)(C(C)(C)C(O)CC2)CC4)CC1 |
| 14 | β2AR; CysLT | 6rz4A | O=C(NCCc1ccc(O)cc1)C1(C)CC2C(C)(CCC3(CC)C4(C)C(C(=O)C=C23)C2(C)C(C)(C(C)(C)C(O)CC2)CC4)CC1 |
| 15 | BambL | 6zfcA | O=C(NC(C(=O)N)Cc1nnn(C2C(O)C(O)C(OC3C(OC4C(O)C(O)C(O)C(C)O4)C(O)C(O)C(CO)O3)C(CO)O2)c1)C(NC(=O)C)Cc1nnn(C2C(O)C(O)C(OC3C(O)C(O)C(O)C(CO)O3)C(CO)O2)c1 |

| | | | |
|---|---|---|---|
| 16 | BCATm | 5hneA | Brc1sc(C(=O)N2CC(Oc3c(C(=O)NC)ccc(-c4c(N=S(=O)([O-])C)cccc4)c3)CC2)cc1 |
| 17 | Bcl-xL | 3wixA | Clc1ccc(C(=O)n2c(C)c(CC(=O)NC(CC(=O)[O-])Cc3c4c(ccc3)cccc4)c3c2ccc(OC)c3)cc1 |
| 18 | BET | 7laiA | O(CC(COC)n1c(C2=CN(C)C(=O)C(C)=C2)nc2c1ccc(N1CCOCC1)c2)C |
| 19 | β2AR | 5a8eA | S(=O)(=O)(N(C(C)C)C)c1cc(C(=O)NC(CC(=O)N)Cc2ccc(C(C)(C)C)cc2)c(Sc2ccc(OC)cc2)cc1 |
| 20 | BRD2-BD2 | 5xhkA | O=C(Nc1cc2C3(C(=O)Nc2cc1)C(C(=O)OCCCC)CC1[NH+]3CSC1)C=Cc1cc(OC)c(OC)c(OC)c1 |
| 21 | BRD4 | 5t35A | O=C(NC)C(NC(=O)c1c(OC(C)C)cc(C2=CN(C)C(=O)C(C)=C2)cc1)CCCCCNC(=O)COc1c2C(=O)N(C(=O)c2ccc1)C1C(=O)NC(=O)CC1 |
| 22 | BRD4 | 5t35D | O=C(NC(C(C)(C)C)C(=O)N1C(C(=O)NCc2ccc(-c3c(C)ncs3)cc2)CC(O)C1)CCC1CCN(c2nc(N(CCC(=O)NC3CC3)C)nc(N(Cc3c(C)n(C)nc3C)C)n2)CC1 |
| 23 | BRDT-BD1 | 7uboA | O=C(NC)C(NC(=O)c1ncc(-c2ccccc2)cc1)CC1CCN(C(=O)Cc2cc3C(C)=CC(=O)Nc3cc2)CC1 |
| 24 | BRDT-BD1, BRDT-BD2 | 7l9aA | O=C(NC)c1ccc(C(=O)Nc2c(C)ccc(NC(=O)c3cc4c(-c5c(C)cc(N)cc5)nn(C)c4cc3)c2)cc1 |

| | | | |
|---|---|---|---|
| 25 | BTK | 4y93A | O=C(C=C)N1C2CN(c3nc(NC)nc(Nc4n[nH]cc4)n3)C(C1)C2 |
| 26 | c-MET | 8ansA | FC(F)c1c(C(=O)N(C)C2c3c(C)ccc(-c4cc5[nH]nc(C)c5cc4)c3CCC2)nccc1 |
| 27 | c-Src | 3el7A | O=C(N)C(NC(=O)CN(C(=O)CCc1ccc(O)cc1)CC1CC1)Cc1cnccc1 |
| 28 | CA-12 | 4ww8A | S(=O)(=O)(N)c1ccc(C(=O)NCc2nnn(C3C(O)C(O)C(O)C(CO)O3)c2)cc1 |
| 29 | CBX2 | 5epkA | Fc1ccc(CC(NC(=O)c2cc3NC(=O)Nc3cc2)C(=O)NC(C(=O)N)Cc2ccc(-c3ccccc3)cc2)cc1 |
| 30 | CDK9, CDK7 | 1ua2A | Clc1c(-c2c3n(nc2)CC(C)(C)C3)cc(NC(=O)Cc2nc(C#N)ccc2)nc1 |
| 31 | CDK9, CDK7 | 3blrA | Clc1c(-c2c3n(nc2)CC(C)(C)C3)cc(NC(=O)Cc2nc(C#N)ccc2)nc1 |
| 32 | CREBBP bromodomain | 4tqnA | S(C(=O)NCCNC(=O)c1c2NC(C)CC(=O)Nc2ccc1)C(S)c1ccccc1 |
| 33 | CREBBP bromodomain | 4yk0A | S(C(S)c1ccccc1)CC(=O)NCCNC(=O)c1c2NC(C)CC(=O)Nc2ccc1 |
| 34 | cyclophilin D (CypD) | 7thdA | O=C([O-])C(C(=O)[O-])Cc1ccc(-c2ccc(CC3C(=O)NCc4c(cccc4)CC(=O)NC(C(=O)NCCOCC[NH3+])CCNC(=O)C=CC(=O)N4CC(Cc5ccccc5)(C(=O)N3)CCC4)cc2)cc1 |

| | | | |
|---|---|---|---|
| 35 | DDP-4 | 2bucA | FC(F)(F)c1n2c(nn1)C1N(C(=O)CC([NH3+])Cc3c(F)cc(F)c(F)c3)C(C2)CC1 |
| 36 | DDR1 | 4ckrA | Brc1c2C(C(=O)N(CC(=O)NCC(F)(F)F)c2ccc1)C1CCN(C(=O)c2nc3c([nH]nc3)cc2)CC1 |
| 37 | MRS; IRS; MetAP; UppS; DHFR | 3nz6X | O=C(Nc1nc(C)ccc1)Cc1c(C(=O)[O-])[nH]c2c1cccc2 |
| 38 | folate receptor (FR) on live regular HeLa cell, EGFR on live A431 cells | 5u8lA | S(CCC(NC(=O)C(=NOC)c1nc(N)sn1)C(=O)NC(C(=O)O)Cc1c2c(sc1)cccc2)C |
| 39 | sEH, c-KIT, ERα | 7te7A | O(C(c1c(C)cccc1)c1c(C)cccc1)C(c1c(C)cccc1)c1c(C)cccc1 |
| 40 | Fibroblast activation protein (FAP) | 1z68A | FC1(F)CN(C(=O)CNC(=O)c2c3c(c(NC(=O)CCC(=O)NC(C(=O)NC(C(=O)NCCNC(=O)CCC(C(=O)O)[NH+]4CC[NH+](CC(=O)O)CC[NH+](CC(=O)O)CC[NH+](CC(=O)O)CC4)Cc4ccccc4)CCCNC(N(C)C)=N)ccc3)ncc2)C(C#N)C1 |
| 41 | FKBP | 5d75A | O=C(O)COc1ccc(CC2C(=O)N3C(C(=O)N(C)C(CC(C)C)C(=O)N(C)CCC=CC(=O)OCC(C)(C)C(=O)C(=O)N4C(C(=O)OC(CCc5cc(OC)c(OC)cc5)c5cc(NC(=O)CCC(=O)NC(c6ccccc6)C(=O)N2)ccc5)CCCC4)CCC3)cc1 |

| | | | |
|---|---|---|---|
| 42 | folate receptor (FR) on live regular HeLa cell, EGFR on live A431 cells | 4kmzA | F[F+2]1C=C2C(=O)C(C(=O)NC(C(=O)NC(C(=O)O)(C)c3c4c(ccc3)cccc4)Cc3c4c(sc3)cccc4)=C[NH+]3C(C)CCC(=C23)C1 |
| 43 | Furin | 7lcuA | Clc1cc(Cl)cc(-c2nc(Oc3cnc(N4CC[NH+](C)CC4)nc3)cc(C[NH+]3CCC(CC(=O)[O-])CC3)c2)c1 |
| 44 | GID4 | 7u3jA | O=C(NC)C(NC(=O)C(NC(=O)C[NH2+]Cc1c(OC)cc(OC)cc1)c1sccc1)CCc1ccccc1 |
| 45 | GSK-3β | 1q5kA | O=C(NCC1CCC(C(=O)NC)CC1)c1c(-c2ccc(OC)cc2)ocn1 |
| 46 | HAO1 | 6w4cA | O=C([O-])c1n[nH]c2c1cc(NCc1c(O)c(-c3onc(N(C)C)n3)ccc1)cc2 |
| 47 | HDAC6 | 5ef7A | O=C(NO)C=Cc1ccc(C[NH2+]CCc2c(C)[nH]c3c2cc(C(=O)N(CC(=O)NCCc2c4c([nH]c2)cccc4)C)cc3)cc1 |
| 48 | HSA | 1bj5A | O=C(OC1CCN(C(=O)c2ccc(-c3[nH]c4c(n3)cccc4)cc2)CC1)Nc1cc(C(=O)NCCOCCOCC[NH3+])ccc1 |
| 49 | Hsp70 | 4po2A | O=C(NC)C(NC(=O)c1ccc(NC(=O)CCC(=O)c2ccc(-c3ccccc3)cc2)cc1)CNC(=O)c1c(O)ccc(NC(=O)c2c(C)onc2)c1 |
| 50 | hTNF-α | 6ooyA | O=C1C(O)=C(c2ccc(O)cc2)Oc2c1c(O)cc(O)c2 |

| | | | |
|---|---|---|---|
| 51 | IDE | 4lteA | O=C(c1ccc(CC2C(=O)NCCCCC(C(=O)N)NC(=O)C=CC(=O)NC(CCCC[NH3+])C(=O)NC(CC3CCCCC3)C(=O)N2)cc1)c1ccccc1 |
| 52 | IDO1 | 5ek4A | Brc1cc2C(NC(=O)c3c4c(cn(COP(=O)(O)O)c4)ccc3)CC3(Oc2cc1)CCN(C(=O)OCC)CC3 |
| 53 | IL-2 | 1m48A | O=C([O-])C(NC(=O)CCn1c(C)cc2c1cccc2)Cc1ccc(C#CCCC)cc1 |
| 54 | IL-17A | 5vb9A | Fc1cc2OCC3(C(C(NC(=O)C4=C(CC)NON4)C(=O)Nc4cnc(-c5c(C)[nH]nc5C)cc4)c2cc1)CC3 |
| 55 | InhA NADH | 5g0sA | O=C(NC)CN1C(=O)C(Cc2ccccc2)N(C(=O)CC2CCC(NC(=O)Cc3c(C)c4c(s3)cccc4)CC2)CC1 |
| 56 | Insulin Receptor | 2z8cA | O=C1[NH+]2C(=Nc3c1cccc3)c1[nH]c3c(c1CC2)cccc3 |
| 57 | Integrin LFA-1 | 1rd4A | Brc1ccc(C(Nc2nc(NC3(C(=O)NCC)Cc4c(cccc4)C3)nc(N3C4CN(C(=O)c5cc(C(F)(F)F)ccc5)C(C3)C4)n2)C)cc1 |
| 58 | MRS; IRS; MetAP; UppS; DHFR | 1jzqA | Clc1cc(Cl)cc(N2CCN(C(=O)c3cc(C(C)(C)C)cc(C[NH2+]Cc4cc(C(=O)NC)ccc4)c3)CC2)c1 |
| 59 | JAK3 | 5ttuA | ClCC(=O)Nc1cc(CNc2nc(NC)nc(Nc3n[nH]c(C(C)(C)C)c3)n2)ccc1 |

| | | | |
|---|---|---|---|
| 60 | JNK1 | 4l7fA | Fc1c(F)ccc(C(=O)C=CC(=O)NCCOCCOCCNC(=O)c2c(Oc3ccccc3)nccc2)c1 |
| 61 | KMO | 4j36B | Clc1c(C(=O)N(C)C2C(C(=O)O)C=C(F)C(Cl)=C2)nc(N2Cc3c(ccc(OC)c3)C2)s1 |
| 62 | KRAS mutant G12C | 6n2kA | Clc1c2c(N3Cc4nc(OCC56[NH+](CCC5)CCC6)cc(N5CC(CC#N)N(C(=O)C(F)=C)CC5)c4CC3)cccc2ccc1 |
| 63 | LC3B | 2zjdA | O=C([O-])CC(n1c(-c2cc(O)c(O)c(C=O)c2)[nH+]c2c1ccc(C(=O)NC)c2)c1cc(Oc2ccccc2)ccc1 |
| 64 | LpxA | 5dg3B | Clc1c(SCC(=O)N(CC(=O)NC)Cc2ccc(-n3ncnc3)cc2)cccc1 |
| 65 | MAP2K6 | 8p7jA | Clc1c(Cl)c(C(=O)C(=C)CC)ccc1OCC(=O)O |
| 66 | Mcl-1; Bcl-2; Bcl-XL | 6oqdA | Clc1cc2c(-c3c(OCC(=O)NC4(C(=O)NC(C(=O)N)CNC(=O)C(NC5=NS(=O)(=O)c6c5cccc6)C(C)C)CC4)cccc3)n[nH]c2cc1 |
| 67 | MDM2, hTEAD4 | 4j3eA | Clc1cc2[nH]cc(C(N(C(=O)CCC(=O)OC(C)(C)C)Cc3nnn(CC(=O)Nc4c(C)cc(C)cc4)c3)C(=O)NC3CCCCC3)c2cc1 |
| 68 | MEK2 | 1s9iA | Fc1c(-c2ccc(C(=O)C(=C)CC(=O)N3C(C(=O)N)CC(Oc4ccccc4)C3)cc2)ccc(F)c1 |
| 69 | Mer, Axl | 7aw2A | Clc1c(-c2ccc(-c3oc(N(CC4=Cn5c(ncc5)C=C4)C)nn3)cc2)cccc1 |

| | | | |
|---|---|---|---|
| 70 | Mer, Axl | 7aazA | O=C(NC1C(OCc2ccc(-c3ccc(C[NH+]4CC[NH+](C)CC4)cc3)cc2)CCC1)c1c(N)ncc(-c2cn(C)nc2)c1 |
| 71 | MRS; IRS; MetAP; UppS; DHFR | 3d27A | Clc1c2n(Cc3[nH]c(C)nn3)c(C(Cc3cc4OCOc4cc3)C)nc2ccc1 |
| 72 | MIF muclesse | 6c5fB | Fc1cc2C(CCc3cc(OC)c(OC)cc3)OC(=O)C3N(C(=O)C(=O)C(C)(C)COC(=O)C=CCCN(C)C(=O)C4N(C(=O)C(COC(C)(C)C)N(C)C(=O)C(c5ccccc5)NC(=O)C(C)N(C)C(=O)COc(c1)c2)CCC4)CCCC3 |
| 73 | MMP-3 | 1g49B | S(=O)(=O)([O-])c1c(C=Cc2c(S(=O)(=O)[O-])cc(NC(=O)C)cc2)ccc(NC(=S)NCc2ccc(CNC(=O)CCc3ccc(OCC4CCCCC4)cc3)cc2)c1 |
| 74 | Mpro | 8skhA | FC(F)(F)C1=CC(C2C(C(=O)NC(C(=O)NC)Cc3ccccc3)CN(C(=O)C(NC(=O)C=C)CC(=C)C)C2)CC=C1 |
| 75 | Mps1 | 5z33A | FC(F)(F)Oc1cc(c(OC)cc1)-c1c2nc(C(=O)NC(C(=O)NC)Cc3c4c([nH]c3)cccc4)ccc2cnc1 |
| 76 | MRS; IRS; MetAP; UppS; DHFR | 7wpiA | Brc1c(O)cc(-c2n(C(CNC(=O)COc3c(C(C)(C)C)cccc3)c3cc(OC)c(OC)cc3)c3c(n2)cc(C(=O)NC)cc3)cc1 |
| 77 | Naa50 | 7ojuA | O=C(NC)C(NC(=O)C1NC(=O)CCC1)CC1CCN(C(=O)c2n(C)nc(C(C)(C)C)c2)CC1 |

| | | | |
|---|---|---|---|
| 78 | NK3 | 7ukzA | O=C(NC)c1cc2nc(n(CC)c2cc1)-c1c(-c2c(C)cc(C)cc2)cccc1 |
| 79 | NMP-2 | 1qibA | O=C(NCCNC(=O)CCc1c(C)n(C2=NC(C)=CC(=O)N2)nc1C)CC(=O)C=CCCCCNC(=O)c1cc(O)cnc1 |
| 80 | OGT | 4n39A | O=C(NC)CC(NC(=O)c1cc(-c2ccc(-c3occc3)cc2)cnc1)c1ccc(-c2ccccc2)cc1 |
| 81 | OXA-48 Carbapenemase | 6pt5A | S(=O)(=O)(O)C1CCN(c2nc(N3CCN(c4c(OCC)cccc4)CC3)ncn2)CC1 |
| 82 | p38 MAP | 1cm8A | O=C(N)C(NC(=O)c1ccc(CNC(=O)c2c(N)n(-c3ccccc3)nc2)cc1)CCC1CCCCC1 |
| 83 | p38α MAP | 7pvuB | O=C(NCc1ccc(C(=O)NCCCC2CCCCC2)cc1)c1c(N)n(-c2ccccc2)nc1 |
| 84 | p38α MAP | 5larA | O=C(N)C(NC(=O)c1ccc(CNC(=O)c2c(N)n(-c3ccccc3)nc2)cc1)CCC1CCCCC1 |
| 85 | p300/CBP | 7ss8A | Clc1ccc(C2(C(=O)N3C(C(=O)NCc4cc(C(=O)NC)ccc4)CCC3)CCCC2)cc1 |
| 86 | PAD4 | 1wd8A | O=C(N1CC([NH3+])CCC1)c1cc2nc(n(C)c2cc1)-c1n(C)c2ncccc2c1 |
| 87 | PAK4, 2-epimerase | 8ahgA | O=C(NC)c1c2c(C(C)C)n[nH]c2cnc1 |
| 88 | PAK4, 2-epimerase | 8aheA | S(=O)(=O)(NC)c1c(C)n(-c2ccccc2)nc1 |

| | | | |
|---|---|---|---|
| 89 | PAR2 | 5nddA | O=C(NC1c2c(C(C(=O)NC)CC1)cccc2)C(NC(=O)c1nnn(Cc2ccccc2)c1)CC1CCCCC1 |
| 90 | PARP1 | 7kkpA | O=C1c2c(O)cc(O)cc2OC(c2cc(O)c(O)cc2)=C1 |
| 91 | PARP10 | 5lx6A | O=C(NC)C(NC(=O)c1cc2c(C(=O)CCC2)cc1)CNC(=O)CCC=1C(=O)Nc2c(N=1)cccc2 |
| 92 | PARP1; PARP15; SIRT6 | 7pw3A | O=C(N)C(NC(=O)c1cc(C)ccc1)CNC(=O)c1cc(C(=O)N)c(C(=O)N)cc1 |
| 93 | PCAF | 3uvwA | Clc1c(Cl)c(C(=O)C(=C)CC)ccc1OCC(=O)NC(C(=O)N)COCCCCCCCCCC |
| 94 | PDE12 | 4z0vA | O=C(N1C(CO)C(c2ccccc2)OCC1)c1cc2[n+](C)c(-c3c4c([nH]c3)cc(C#N)cc4)[nH]c2cc1 |
| 95 | PI3K | 3apcA | O=C(O)C(NC(=O)CCC(NC(=O)C1=C(C)n2nc(C)cc2N=C1)C(=O)Nc1cc(OC)c(OC)c(OC)c1)CCC(=O)N |
| 96 | PI3K (WT/H1047R) | 4yknA | S(=O)(=O)(N=C1SC(C)=NN1)c1ccc(NCc2cc(C(=O)[O-])cc(-c3cnc(OC)cc3)c2)cc1 |
| 97 | Pin1 | 2xpbA | ClCC(=O)N1CCN(c2nc(NC(c3cc(S(=O)(=O)C)ccc3)c3ccccc3)nc(NC)n2)CC1 |
| 98 | PqsE Thioesterase | 7kgwA | Clc1cc(OC2CN(C(OC(C)(C)C)=O)CCCC2)cc(NC(=O)c2c(C(=O)O)cccc2)c1 |
| 99 | prolyl hydroxylases | 5c5tA | O=C([O-])c1ncn(C2=NC(=O)NC(NCc3ccc(-c4ccccc4)cc3)=N2)c1 |

| | | | |
|---|---|---|---|
| 100 | prion protein (PrP) | 4ma7A | O=C(NCCOCCOCC=N)C1NC(=O)C2CN(C(=O)CCCN(C)C(=O)C3N(C(=O)C=CC(=O)NCCC1)CCC3)CC2 |
| 101 | PSMA | 1z8lA | O=[N+]([O-])c1c(N2CCCC2)ccc(C(=O)NCC(NC(=O)CC23CC4CC(C2)CC(C3)C4)C(=O)NC)c1 |
| 102 | RIP2 | 5j79A | Clc1c(O)cc(NCc2cc(C(=O)NCCOC)cc(-c3cc4[nH+]ccc(Nc5cc(O)c(Cl)cc5)c4cc3)c2)cc1 |
| 103 | ROCK2 | 6ed6A | O=C(N1C(c2ccccc2)C[NH2+]CCC1)c1cc2c(cncc2)cc1 |
| 104 | RORγ Reporter (Gal4) | 6q2wA | FC(F)(F)C(O)(C(F)(F)F)c1ccc(CN(c2c(F)c(NCC3C(O)C[NH+](CC(=O)N)CC3)ncn2)C2CC2)cc1 |
| 105 | RSA; BSA; HSA | 6wuwA | FC(F)(F)c1cc(CC(=O)NC(C(=O)NC)CNC(=O)Cc2cc([N+](=O)[O-])ccc2)ccc1 |
| 106 | RSV N protein | 4uccA | Brc1sc(CC2NC(=O)C(Cc3ccccc3)NC(=O)C3N(C(=O)Cn4nnc(c4)CC(C(=O)N)NC(=O)C(C(CC)C)CNC(=O)C(Cc4cc(OC)ccc4)NC(=O)c4cc(NC2)ccc4)CCC3)cc1 |
| 107 | sEH | 1zd4A | FC(F)(F)c1c(CNC(=O)C2CC(Nc3nc(NC)nc(N4CC[NH+](C)CC4)n3)CCC2)cccc1 |
| 108 | sEH, c-KIT, ERα | 5mwaA | Clc1c(Sc2ccc(NC(=O)C3CCN(C(=O)c4ccc(F)cc4)CC3)cc2)cccc1 |

| | | | |
|---|---|---|---|
| 109 | Sirt3 | 4fvtA | O=[N+]([O-])c1c(C(O)C)cc(OC)c(OCCCC(=O)NC2c3c(cc(OCC(=O)NCCCC)cc3)CCc3c2cccc3)c1 |
| 110 | SIRT3 | 4jsrA | O=C(N(CCCN1c2c(cccc2)CCc2c1cccc2)C)CCC(C)(c1ccc(O)cc1)c1ccc(O)cc1 |
| 111 | SIRT5 | 6ljmA | O=C(OC(C)(C)C)NC(CC(=O)NCCCCCC(=O)N(CCCN1c2c(cccc2)CCc2c1cccc2)C)Cc1ccccc1 |
| 112 | SOS1 | 5oviA | O=C(OC(C)(C)C)N(Cc1c(-c2cc(C(NC3=NNC(=O)c4c3cc(N3CC[NH+](C)CC3)nc4)C)sc2)cccc1)C |
| 113 | Src kinase | 2srcA | Fc1ccc(CC2C(=O)NC(Cc3ccc([N+](=O)[O-])cc3)C(=O)NCCCC(C(=O)[O-])NC(=O)C=CC(=O)NC(CC3CCCCC3)C(=O)N2)cc1 |
| 114 | STING | 8gt6A | S(C)c1c2n(CC=CCn3c(NC(=O)c4c(CC)nc(C)s4)nc4c3c(OCCC[NH+]3CC[NH+](C5CC5)CC3)cc(C(=O)N)c4)c(NC(=O)c3n(CC)nc(C)c3)nc2cc(C(=O)N)c1 |
| 115 | Streptavidin | 1lcvA | O=C(N(Cc1ccc(C(=O)N(Cc2sc(C[NH+](Cc3ccc(C(=O)N)cc3)C)cc2)C)cc1)CC1CCOCC1)c1ccc(C[NH2+]CC2CCOCC2)cc1 |
| 116 | TAK1 | 4l53A | FC(F)Oc1c(CN(C(=O)c2cc(C(=O)N3CCCC3)[nH]c2)CC(=O)NC)cccc1 |

| | | | |
|---|---|---|---|
| 117 | Tankyrase 1 | 6qxuA | O=C(NCCc1onc(-c2ncccc2)n1)c1c(N2C(=O)NC(=O)CC2)cccc1 |
| 118 | YAP-Binding domain of TEAD1 | 8cuhA | Clc1c(C[NH2+]C(C(=O)NC(C(=O)N)c2ccccc2)C2Cc3c(cccc3)C2)c(O)cc(Cl)c1 |
| 119 | Thrombin | 1dwbH | O=C(N(C)C1c2c(OCC1)cccc2)NC(C(=O)N1CC(C(=O)NC)C(c2ccc(OC)cc2)C1)Cc1c(CNC(=[NH2+])N)cccc1 |
| 120 | TNKS1; HSA | 3uh2A | FC1(F)Oc2c(O1)ccc(C(=O)NCc1cc(C(=O)N)cc(CNC(=O)CCc3cc(OC)c(OC)c(OC)c3)c1)c2 |
| 121 | tankyrase 1 (TNKS1) | 4i9iA | O=C(NC)C(NC(=O)c1cnc2c(c1)cccc2)CNC(=O)CCN1C(=O)NC(=O)c2c1cccc2 |
| 122 | TRK | 5wr7A | Fc1cnc2c(c1)C1N(C3=NC=4[NH+](N=CC=4C(=O)NC(C)CC2)C=C3)CCC1 |
| 123 | TRKA | 6iqnA | Fc1cc2c(OCCC(C)NC=3C(C#N)=CNC=4C=3NC(N3C2CCC3)=CC=4)cc1 |
| 124 | Trypsin | 2staA | S(=O)(=O)(NC(C(=O)NC(C(=O)NCc1ccc(C(=[NH2+])N)cc1)C)CNC(=O)Cc1cc2c(cc1)cccc2)c1ccccc1 |
| 125 | MRS; IRS; MetAP; UppS; DHFR | 5kh5B | Clc1c(Cl)cc(C(=O)NCc2ccc(C(C)(C)C)cc2)c(C(=O)NC(C(=O)NC)C(C)(C)C)c1 |
| 126 | VCP/p97 | 4kdiA | SC1=[NH+]c2c(C(=O)N1)ccc(C(=O)Nc1ccc(C=CC(=O)N)cc1)c2 |
| 127 | Wip1 | 1a6qA | Clc1cc(NCc2sc(C(=O)NC(C(=O)NCCOC)CC3CCCCC3)cc2)ccc1 |
| 128 | Z α1-antitrypsin | 2qugA | O=C(NC(C(O)c1ccccc1)CCC)c1cc2[nH]ccc2cc1 |
